# Crystal structure of the MBD domain of MBD3 in complex with methylated CG DNA

**DOI:** 10.1101/338061

**Authors:** Ke Liu, Ming Lei, Bing Gan, Harry Cheng, Yanjun Li, Jinrong Min

## Abstract

MBD3 is a core subunit of the Mi-2/NuRD complex, and has been previously reported to lack methyl-CpG binding ability. However, recent reports show that MBD3 recognizes both mCG and hmCG DNA with a preference for hmCG, and is required for the normal expression of hmCG marked genes in ES cells. Nevertheless, it is not clear how MBD3 recognizes the methylated DNA. In this study, we carried out structural analysis coupled with isothermal titration calorimetry (ITC) binding assay and mutagenesis studies to address the structural basis for the mCG DNA binding ability of the MBD3 MBD domain. We found that the MBD3 MBD domain prefers binding mCG over hmCG through the conserved arginine fingers, and this MBD domain as well as other mCG binding MBD domains can recognize the mCG duplex without orientation selectivity. Furthermore, we found that the tyrosine-to-phenylalanine substitution at Phe34 of MBD3 is responsible for its weaker mCG DNA binding ability compared to other mCG binding MBD domains. In summary, our study demonstrates that the MBD3 MBD domain is a mCG binder, and also illustrates its binding mechanism to the methylated CG DNA.

## INTRODUCTION

5-methylcytosine (5mC), a product of DNA cytosine methylation, is an epigenetic mark in both animals and plants that is essential for various biological processes, such as X-chromosome inactivation, genomic imprinting, transposon silencing, and transcriptional repression (1). DNA cytosine methylation mediates transcriptional regulation through binding to a family of proteins that contain an approximately 70 residues methyl-CpG-binding domain (MBD). Aberrant DNA methylation, such as hypermethylation of tumor suppressor genes, has been linked to various human diseases, including cancers (2).

The first purified protein complex (MeCP1 complex) with methyl-CpG binding activity was identified in 1989 (3). MeCP2 is the first protein to be cloned with methyl-CpG binding ability (4), and its MBD was shown to bind methyl-CpG DNA directly (5). Based on sequence homology, MBD1 was identified as the methyl-CpG binding protein in the MeCP1 complex (6). Several other MBD-containing proteins have been identified since (4,7–12). The human genome encodes 11 known MBD-containing proteins: including MeCP2, MBD1-6, SETDB1/2 (Histone-lysine N-methyltransferase SETDB1/2) and BAZ2A/2B (Bromodomain adjacent to zinc finger domain protein 2A/2B). Most of the MBD-containing proteins also include other chromatin-associated domains, such as CXXC (CXXC-type zinc finger protein), PHD (plant homeodomains), Bromodomain, and SET domains. The MBD proteins are often associated with histone deactylases, chromatin remodelling complexes and/or histone methyltransferases, which recruit these chromatin modifying activities to DNA-methylated chromatin regions for transcriptional silencing/repression (11).

MBD2 and MBD3, two subunits of the Mi-2 autoantigen (Mi-2)/nucleosome remodelling and histone deacetylase (NuRD) complex (Mi-2/NuRD complex), have been studied extensively regarding their function as chromatin structure regulators (9,13–15). Despite ~70% identity between the MBD domains of MBD3 and its closest homolog MBD2, MBD3 was considered unable to bind mCG DNA (8). It has been reported that both MBD2 and MBD3 are found at CG rich promoters, but MBD2 preferentially binds to methylated CG islands, whereas MBD3 mainly locates at promoters and enhancers of active genes with little cytosine modification (16). On the other hand, electrophoretic mobility shift assays (EMSA) display that MBD3 binds to both mCG and hmCG DNA with a preference for hmCG DNA (17). NMR analysis also supports the binding capacity of the MBD3 MBD domain to mCG (18).

Although recent studies have shown that the MBD domain of MBD3 is able to bind both mCG and hmCG, in this study, we quantitatively characterized the binding ability of the MBD domain of MBD3 to different DNAs, including fully methylated CG (mCG), mCG/TG mismatch and hmCG DNA. We further solved the complex structures of the MBD3 MBD domain with different mCG DNA, including two palindrome mCG DNA and one non-palindrome mCG DNA, together with a MBD1-mCG complex structure. Our structural analysis coupled with binding and mutagenesis data revealed that MBD3 is a mCG binder, and like other mCG binding MBD domains, its MBD domain recognizes the mCG DNA without sequence selectivity outside the mCG dinucleotide.

## RESULTS AND DISCUSSION

### MBD3 MBD domain is able to bind methylated CG DNA via the conserved arginine fingers

The MBD domain of MBD3 was thought to lack mCG binding ability *in vitro* (8,13), and genome-wide distribution analysis by ChIP-seq experiments also failed to show pronounced MBD3 enrichment at methylated islands (19). In contrast, other studies have indicated that MBD3 harbours methylated CG binding ability (18,20), and MBD3 prefers binding to hmCG over to mCG DNA (17). Furthermore, DNA methylation is required for genomic localization of MBD3 *in vivo* (21). In an effort to reconcile the conflicting DNA binding data of MBD3, we compared the binding affinities of the MBD domain of MBD3 to hmCG and mCG DNA, respectively, by ITC assays, and measured only weak binding affinity between the MBD3 MBD domain and hmCG DNA (Fig. 1A and Table 1). On the other hand, MBD3, like MeCP2 and MBD1/2/4, recognized fully methylated CG (mCG) DNA, and similar to previously published results (20), the sequence surrounding the mCG dinucleotide did not affect its binding affinity significantly (Fig. 1A). Notwithstanding strong sequence conservation with the MBD2 MBD domain (Fig. 2A), MBD3 bound to mCG DNA about 5 fold weaker than MBD2 (Table 1). In order to understand the structural basis of mCG binding by MBD3, we determined crystal structures of MBD3 MBD domain in complex with two 12mer palindromic mCG containing DNA, respectively (Table S1).

**Figure 1.**
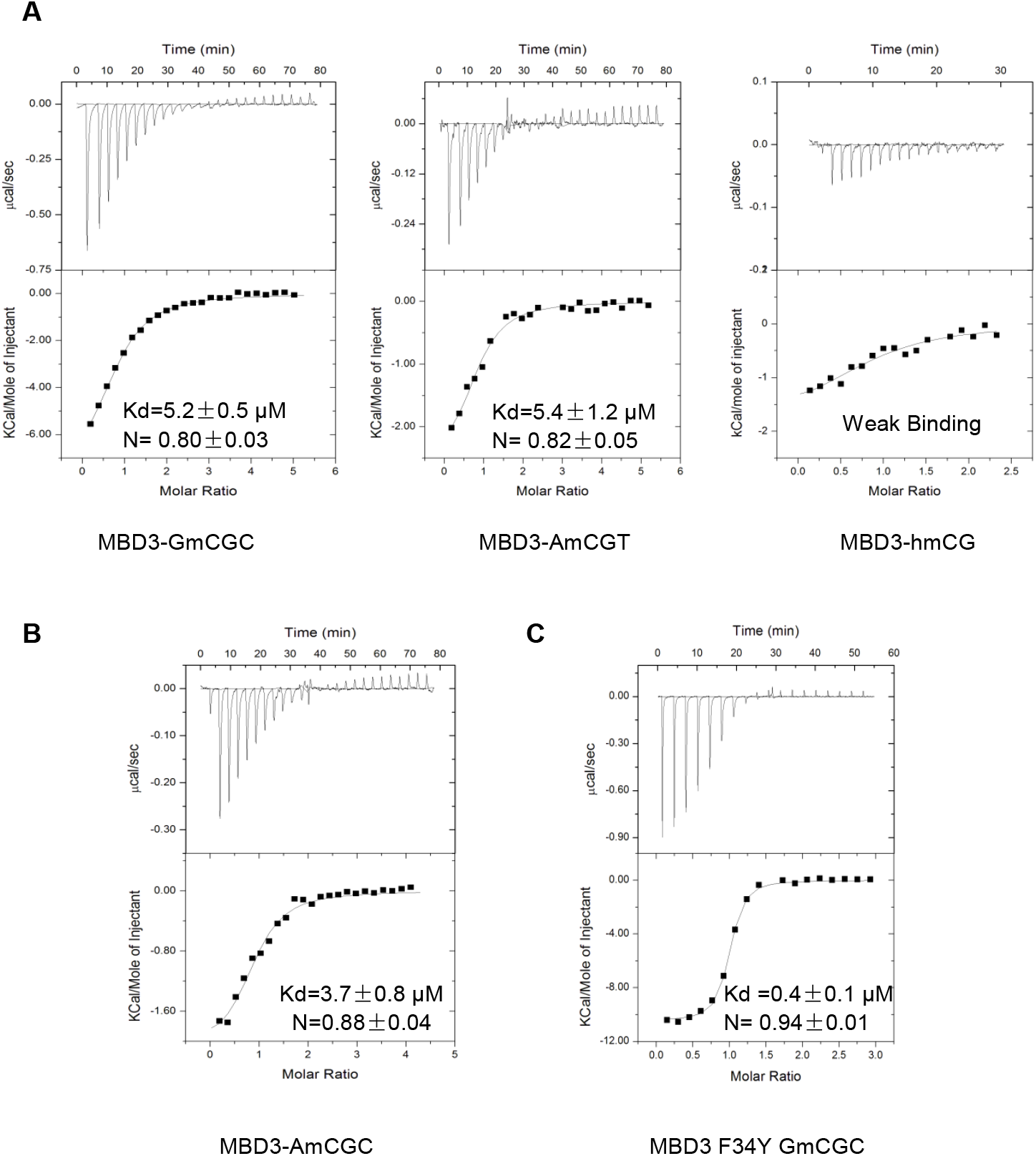
ITC binding curves of the MBD3 WT and F34Y mutant proteins to methylated DNA. GmCGC sequence: 5’-GCCAGmCGCTGGC-3’; AmCGT sequence: 5’-GCCAAmCGTTGGC-3’; hmCG sequence: 5’-GCCAGhmCGCTGGC-3’; AmCGC sequence: upper strand 5’-GCCAAmCGCTGGC-3’, lower strand 5’ -GCCAGmCGTTGGC-3’.

**Figure 2.**
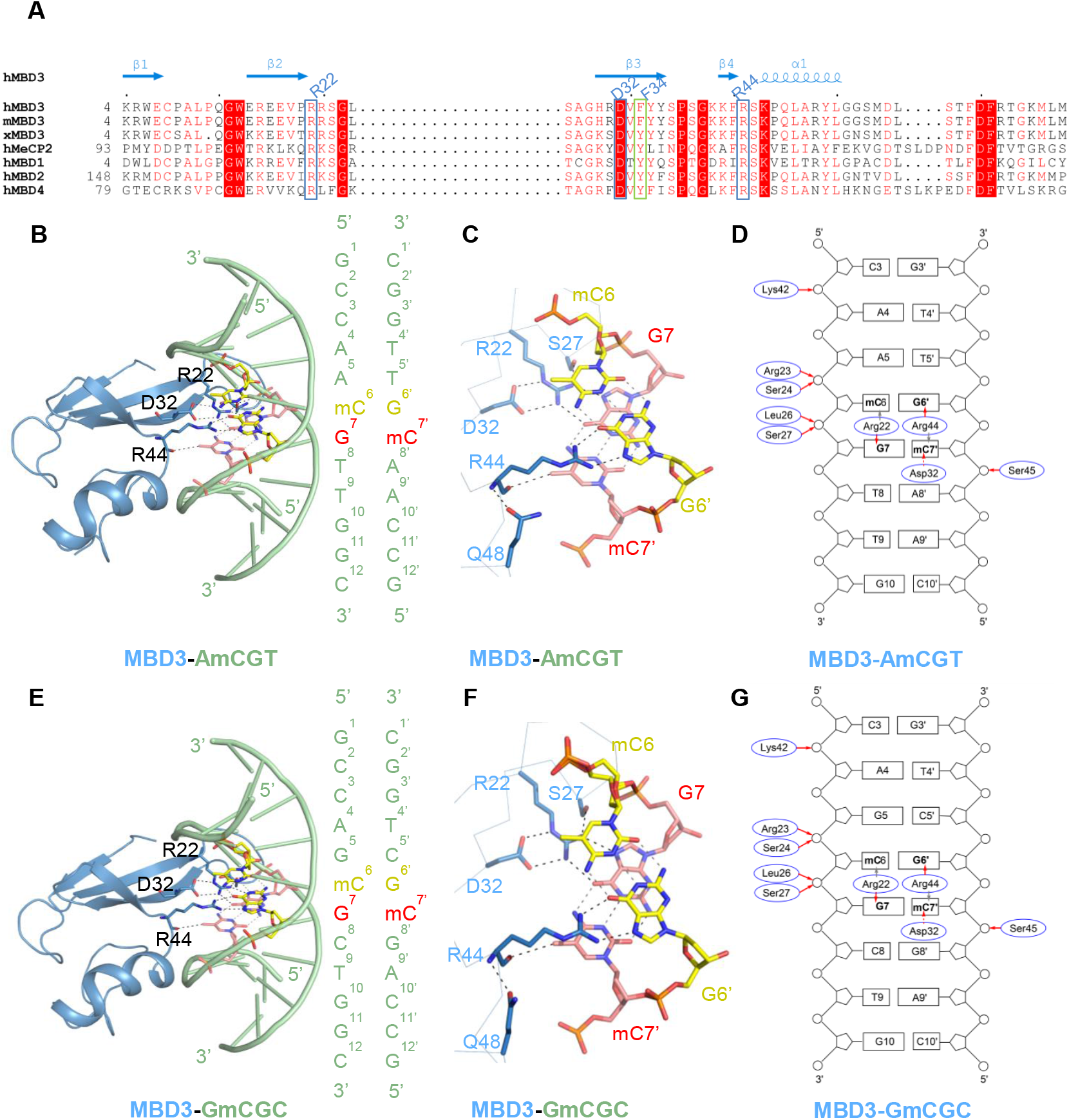
Structural basis of the MBD3 MBD domain in complex with mCG DNA. (A) Structure based sequence alignment of MBD domains. Secondary structure elements and residue numbers involved in DNA binding of MBD3 are shown at the top of the sequence alignment. The alignments was constructed with ClustalW and refined with ESPript. hMBD1, hMBD2, hMBD3, hMBD4 and hMeCP2 represent the MBD domains of human MBD1 (NP_001191065.1), human MBD2 (NP_003918.1), human MBD3 (NP_001268382.1), human MBD4 (NP_001263199.1) and human MeCP2 (NG_007107.2), respectively. xMBD3 and mMBD3 represent the MBD domains of *Xenopus laevis* MBD3 (AAD55389.1) and mouse MBD3 (NP_038623.1), respectively. (B) and (E) Overall structures of the MBD3 MBD domain bound to two different mCG DNA in cartoon representation. The protein and DNA are colored in blue and green, respectively, except the two mC-G base pairs, which are shown in yellow and amaranth sticks, respectively. The mCG dinucleotide binding protein residues are shown in stick models. The dashed lines represent the hydrogen bonds formed between protein and DNA or base pairs. (C) and (F) Detailed interactions between the two mC-G base pairs and the MBD3 MBD domain. Hydrogen bonds formed between protein residues and DNA or base pairs are marked as black dashed lines and grey dashed lines, respectively. (D) and (G) Schematic representation of intermolecular contacts between the MBD3 MBD domain and mCG DNA. The direct interactions and water mediated interactions between protein residues and DNA are shown as red solid and dashed lines, respectively; the grey dashed lines represent stacking interactions.

**Table 1.** Binding affinities (Kds) of the MBD domains of MeCP2 and MBD1-4 with different DNA (μM).

|  | DNA Sequences | MeCP2<br>(aa 80-164) | MBD1<br>(aa 1-77) | MBD2<br>(aa 143-220) | MBD3<br>(aa 1-71) | MBD4<br>(aa 55-152) |
| --- | --- | --- | --- | --- | --- | --- |
| mCG | 5'-GCCAA**mCG**TTGGC-3' | $1.2 \pm 0.2^*$ | $1.0 \pm 0.1^\#$ | $0.9 \pm 0.2^*$ | $5.4 \pm 1.2^\#$ | $0.4 \pm 0.1$ |
| mCG/TG<br>Mismatch | 5'-GTCAA**mCG**TTACG-3'<br>3'-CAGTT**GT**AATGC-5' | $1.2 \pm 0.2$ | $2.2 \pm 0.3$ | $1.6 \pm 0.2$ | $6.5 \pm 1.2$ | $1.5 \pm 0.3$ |
| hmCG | 5'-GCCAG**hmCG**CTGGC-3' | $32 \pm 4$ | WB | $9.7 \pm 2.3$ | WB | WB |

WB: weak binding. For these ITC data, the heat is too little to be fitted accurately. \*: These data are from Reference #20. #: DNA used for co-crystallization in this study.

The two structures are essentially isomorphous. In the respective atomic models, only the mCG dinucleotide duplex made base-specific interactions with MBD3 (Figs. 2B-2G), consistent with the only small effect of flanking base types on binding constants (Fig. 1A) and with structures of homologous MBD-DNA complexes (22–26). Thus, MBD3 coordinates can be aligned with an MBD2 model with an RMSD of 0.9Å over aligned Cα coordinates (20).

In the MBD3-DNA structures, the 4-strand β sheet of MBD3 inserted into the major groove of the mCG DNA near the mCG dinucleotide motif (Figs. 2B and 2E), and each mCG dinucleotide was recognized by an “arginine finger”, i.e., Arg22 on the C terminal tip of the second β-strand and Arg44 on the C terminal tip of the fourth β-strand (Figs. 2B and 2E), forming a stair-shaped interactions pattern (Figs. 2C, 2D and 2F, 2G) (20). Both Arg22 and Arg44 are highly conserved in functioning MBD domains (Fig. 2A). Similar to MBD2, the side chain of Arg22 was fixed by forming a salt bridge with Asp32 and a hydrogen bond with the side chain of Ser27 (Figs. 2C and 2F).

### MBD3 recognizes the two different strands of the DNA duplex without orientation selectivity

Considering that only the mCG dinucleotide makes base-specific interactions, the MBD domain should be able to bind to either strand of the DNA duplex equally well. In order to confirm this, we determined the crystal structure of the MBD domain of MBD3 in complex with a non-palindromic AmCGC DNA. Our ITC binding results revealed that the MBD domain of MBD3 bound to the non-palindromic mCG (AmCGC) as well as to our palindromic mCG DNA in this study (Figs. 1A and 1B). Our complex structure revealed that the electron density map of AmCGC DNA in the MBD3-AmCGC complex was too ambiguous to place it in a certain orientation, or the electron density map of the AmCGC was an averaged map of both orientations of the DNA duplex. Therefore, refining either orientation against the electron density map generated similar statistic results (Figs. 3A-3D and Table S1). Consistently, a stochastic binding of the MBD1 MBD domain to either strand of a mCG DNA duplex has been observed by two-dimensional NMR spectroscopy (27). Hence, we propose that all the mCG binding MBD domains are able to recognize the mCG DNA duplex without orientation selectivity.

**Figure 3.**
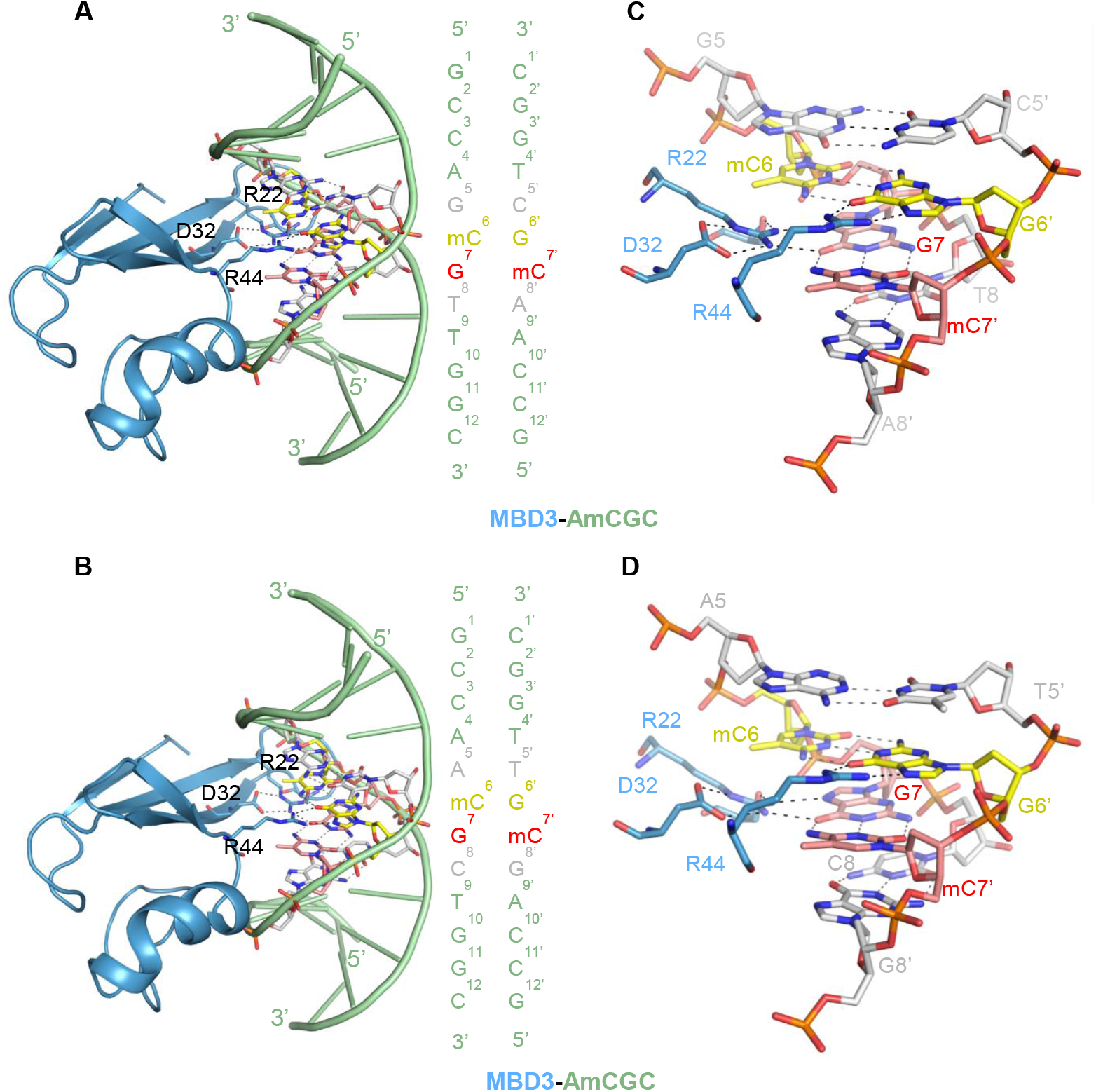
MBD3 recognizes the two different strands of the DNA duplex without orientation selectivity. (A) and (B) The overall view of the MBD3 MBD domain bound to the two different orientations of the DNA duplex. The DNA interacting protein residues are shown as sticks and colored in blue. The central DNA bases are shown as stick models and marked in yellow, red and grey, respectively. (C) and (D) The detailed interactions of the two mC-G base pairs and the MBD3 MBD domain. The protein residues and bases are shown in the same way as (A) and (B). Hydrogen bonds formed between protein residues and DNA are marked as black dashed lines, and hydrogen bonds formed between base pairs are shown as grey dashed lines.

### The two arginine fingers in the MBD domain contribute to the mCG dinucleotide binding not equally

Previously on the basis of the MBD4-mCG/TG complex structure, it has been proposed that the tight recognition of mCG by the first arginine finger (or Arg finger-1) prevents the flipping binding of the MBD4 MBD domain on asymmetric target sequences, such as mCG/TG, mCG/hmCG, and mCG/hmUG (24). Although the mCG duplex in either of the two opposite orientations could be bound equally well by the MBD domain, the two arginine fingers have different binding contexts, possibly making different contributions to the mCG DNA binding. Recently, we have shown that, in the MBD2-mCA complex structure, the first arginine finger R166 of MBD2 is used to recognize the TG dinucleotide because R166 is tightly fixed and cannot tolerate the adenine base of the CA dinucleotide, and the more flexible arginine finger, i.e., the second arginine finger R188, is pushed away by the adenine base of the CA dinucleotide (20). But when we mutated R166 to alanine, the mutated MBD2 could not bind to mCA DNA any more, and the R188A mutant could still bind to mCA (20). If both arginine fingers contributed to the mCA DNA binding equally, when R166 was mutated to alanine, A166 should be able to tolerate the mCA dinucleotide, and R188 would be used to recognize the TG dinucleotide. Now that the R166A mutant does not show any binding to mCA DNA, it implies that the R188 finger contributes much less to the DNA binding. Consistently, our binding results, indeed, showed that the R166A mutant could not bind to mCG DNA, but the R188A mutant still showed comparable binding to mCG DNA (Fig. 4A). Taken together, R166 is a major contributor to the mCG or mCA binding.

**Figure 4.**
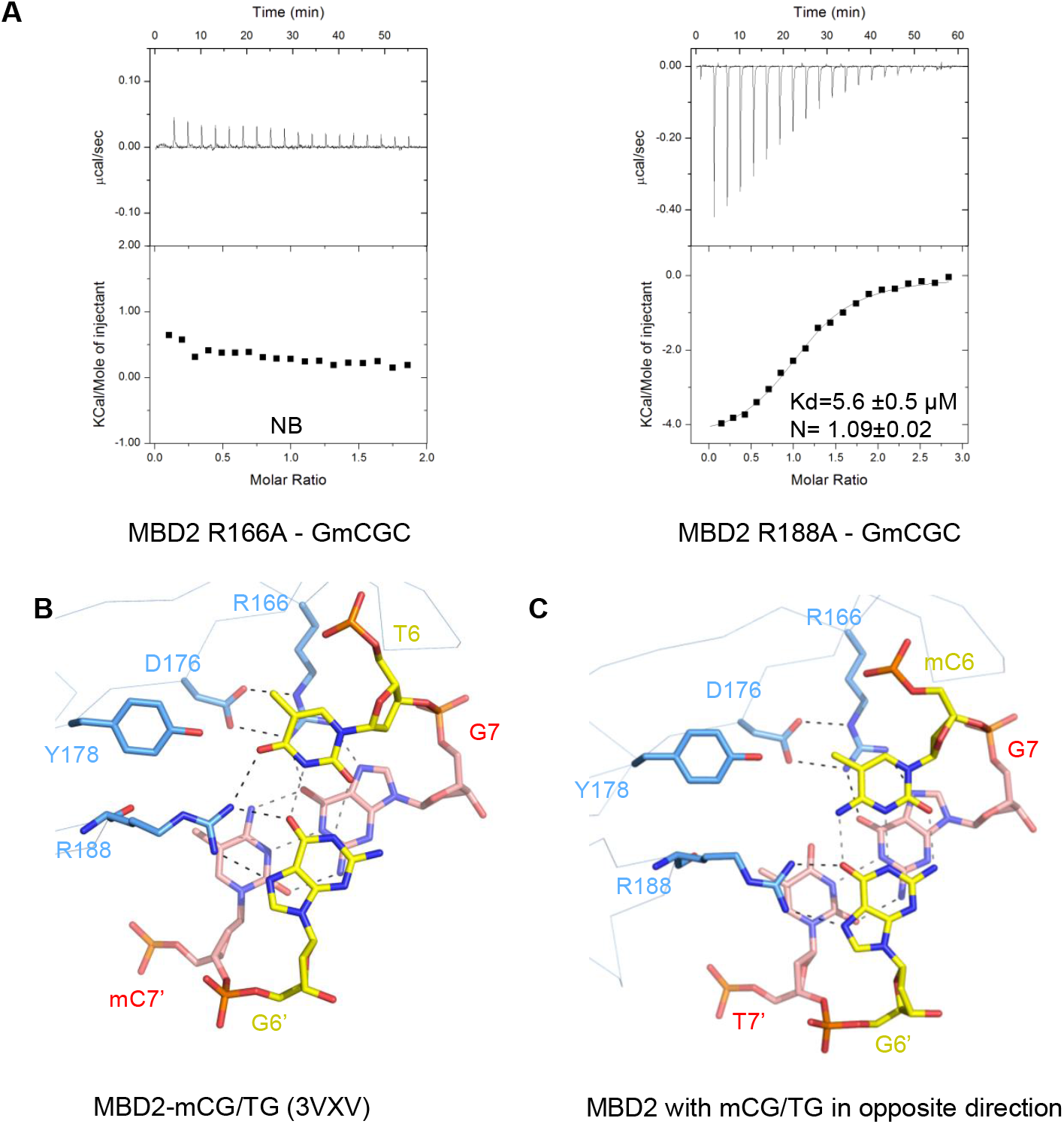
The two arginine fingers in the MBD domain contribute to the mCG dinucleotide binding not equally. (A) The ITC binding curves of the MBD2 mutants to mCG DNA. NB: No detectable binding. (B/C) Models for the MBD2 MBD domain bound to a mCG/TG mismatch DNA based on the published MBD4-mCG/TG complex structure (PDB code: 3VXV) with R166 stacking with the T base (B) and R188 stacking with the T base (C).The mCG/TG interacting protein residues are shown as blue sticks, and the mCG/TG base pairs are shown as yellow and amaranth sticks, respectively. Hydrogen bonds formed between protein residues and DNA are marked as black dashed lines, and hydrogen bonds formed between base pairs are shown as grey dashed lines.

MBD4 is a mismatch-specific DNA N-glycosylase, so it not surprising to find that its MBD domain is able to bind to mCG/TG mismatch DNA (24). In this study, we found that all the mCG DNA binding MBD domains were able to bind mCG/TG as well as mCG/mCG DNA (Table 1), presumably due to the fact that thymine is a mimic of methyl-cytosine. The published MBD4-mCG/TG complex structure showed that it is the first arginine finger of MBD4 to bind the TG dinucleotide through the stair interactions (24). Although the thymine base could only form two hydrogen bonds with its pairing guanine, it forms an extra hydrogen bond with the second arginine finger (Fig. 4B). If the second arginine finger bound to the TG dinucleotide, the thymine base could not form another hydrogen bond with the first arginine finger because the first arginine finger is fixed too rigidly to take a different confirmation to form a hydrogen bond with the thymine base (Fig. 4C).

### The tyrosine-to-phenylalanine substitution at Phe34 of MBD3 is responsible for MBD3’s weaker binding ability

Previously it has been reported that substitutions at two key residues in human/mouse MBD3, i.e., Phe34 and His30, which respectively are tyrosine and lysine/arginine in MBD1/2/4 as well as MeCP2, contribute to the binding inability of human/mouse MBD3 to mCG DNA (13). In contrast, *Xenopus laevis* MBD3 harbours tyrosine and lysine in the corresponding positions and retains mCG DNA binding ability (Fig. 2A) (28). Even though the MBD3 MBD domain was eventually confirmed to bind mCG DNA, it does so less tightly than its paralog MBD2 (Fig. 1A and Table 1) (20). To investigate whether and how the Phe34/His30 substitutions affect its mCG binding ability, we compared the complex structures of MBD3 with those of other MBD domains.

Structural comparisons reveal that the side chain carbonyl group of the tyrosine residue, conserved in MeCP2, MBD1 and MBD2, engages in solvent-mediated interaction with the 4-amino group of the methylated cytosine (Figs. 5A-5C). Even as the corresponding MBD4 Tyr96 side chain points away from the DNA groove (Fig. 5D) and is accommodated in a hydrophobic pocket formed by Val80, Lys82, Ile98 and Lys104 (24), its hydroxyl group engages in solvent mediated hydrogen bonding with the DNA backbone (Fig. 5E). However, MBD3 lacks this solvent-mediated interaction due to the inability of Phe34 in forming such a hydrogen bond, which might explain why MBD3 bound mCG DNA weaker than other mCG binding MBD domains (Figs. 5F and 5G) (20).

**Figure 5.**
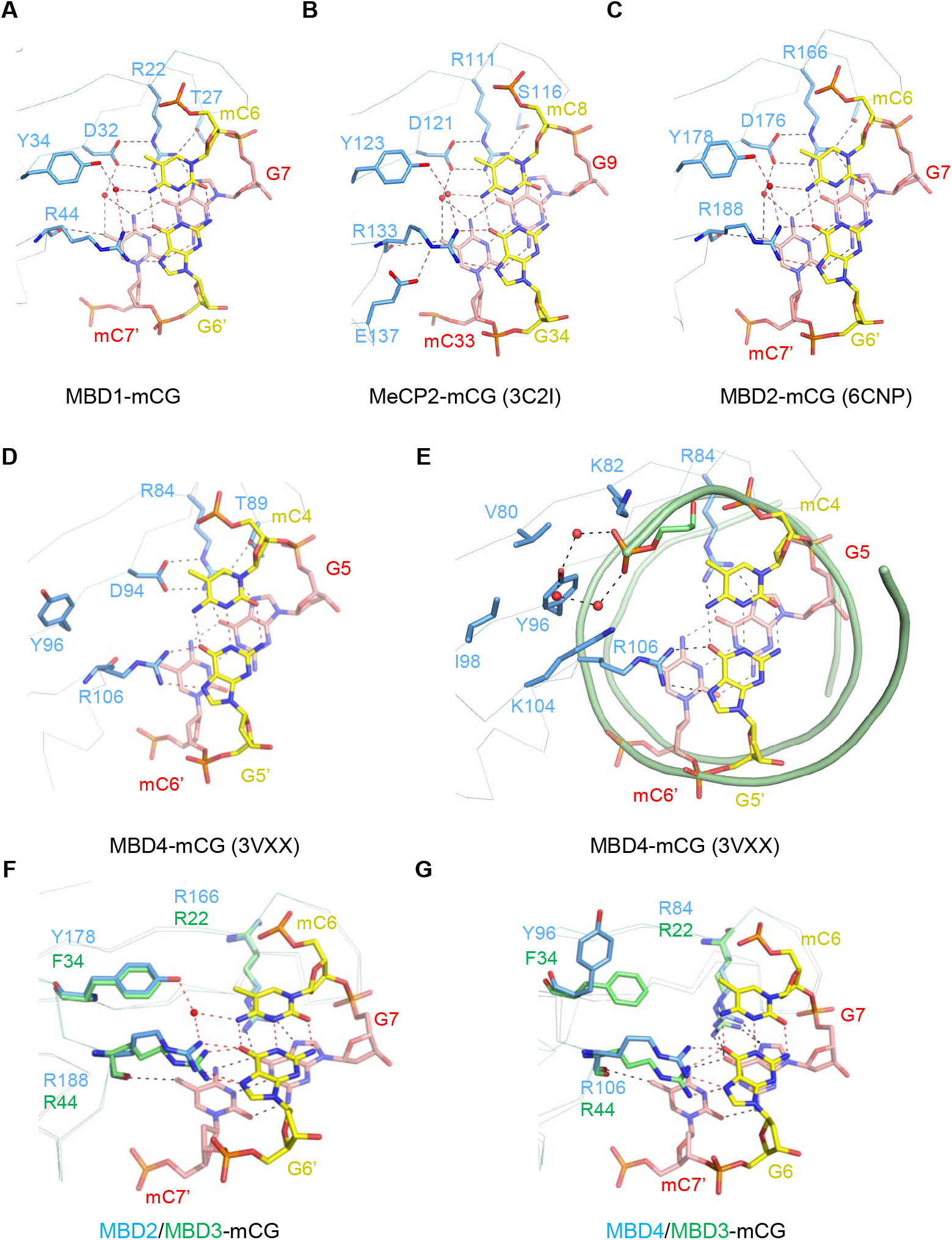
The tyrosine-to-phenylalanine substitution at Phe34 of MBD3 is responsible for MBD3’s weaker mCG DNA binding ability. (A-D) Detailed interactions of the MBD domains of MBD1, MeCP2, MBD2 and MBD4 bound to their corresponding mCG DNA. The mCG dinucleotide DNA and their interacting protein residues are shown as stick models. Water molecules are shown as red balls. (E) Tyr96 of mouse MBD4 is accommodated in a hydrophobic pocket formed by Val80, Lys82, Ile98 and Lys104, and forms water-mediated interactions with the backbone of DNA. The protein residues are shown as blue sticks; mC6’-G5 and mC4-G5’ are shown in red and yellow sticks, respectively. DNA backbone is shown as green cartoon with the Tyr96 interacting phosphate backbone marked as stick models. Water molecules are shown as red balls. Hydrogen bonds formed between Tyr96 and phosphate backbone are marked as black dashed lines. (F) Superimposition of MBD2-mCG and MBD3-mCG structures. (G) Superimposition of MBD3-mCG and MBD4-mCG structures. The protein residues and bases are shown in the same way as in (E).

To further investigate if substitution Phe34 with tyrosine would enhance the mCG binding by MBD3, we made a MBD3 F34Y mutant, and carried out ITC binding studies. Our binding results revealed that the mCG binding affinity of F34Y MBD3 was increased approximately tenfold than that of WT MBD3 (Figs. 1A and 1C). Taken together, crystal structures and affinity data suggest that, although the Phe34 substitution decreases the mCG binding affinity of MBD3, mCG binding by MBD domain does not strictly require the tyrosine residue at this position.

Regarding His30 of MBD3, structural analysis revealed that the His30 of MBD3 is fixed by forming a hydrogen bond with Arg65, and does not form any unfavourable interactions with DNA, consistent with reported MeCP2 and MBD2 structures (Fig. S1). Thus, the His30 residue of MBD3 does not play any significant role in the DNA recognition.

## EXPERIMENTAL PROCEDURES

### Protein Expression and Purification

The MBD domains of human MeCP2 (aa 80-164), MBD1 (aa 1-77), MBD2 (aa 143-220), MBD3 (aa 1-71) and MBD4 (aa 55-152) were subcloned into a modified pET28-MHL expression vector to generate N-terminal His fusion protein for ITC binding assay. For the crystallization, both MBD1 (aa 1-77) and MBD3 (aa 1-71) fragments were subcloned into a pET28-GST-LIC expression vector to generate N-terminal GST fusion protein. The MBD2 (R116A and R188A) and MBD3 (F34Y) mutants were obtained with the MBD2 (aa 143-220) and MBD3 (aa 1-71) pET28-MHL expression constructs, respectively, by Quick Change site-directed mutagenesis (Agilent Technologies). Then these plasmids were transformed into *Escherichia coli BL21 (DE3)-V2R-pRARE2* cell for overexpression. The recombinant protein was induced with 1 mM isopropyl-β-d-thiogalactopyranoside (IPTG) at 14°C. The cells after collection were broken in buffer containing 20 mM Tris • HCl, pH 7.5, 500 mM NaCl, 5 mM βME, 1 mM PMSF. The supernatants were collected after centrifugation and further analyzed by affinity chromatography. For crystallization experiments, the MBD1 and MBD3 proteins were treated with Thrombin to remove the GST tag. The proteins were further purified by anion-exchange column and gel filtration column (GE Healthcare). Finally, the purified protein was concentrated to 10 mg/mL in a buffer containing 20 mM Tris • HCl, pH 7.5, 150 mM NaCl, and 1 mM DTT. For ITC experiment, the pure protein was dissolved in the same buffer without DTT.

### Binding Assays

All DNA oligos used for isothermal titration calorimetry (ITC) measurements were synthesized by IDT (Integrated DNA Technologies, USA). After dissolving in the ITC buffer containing 20 mM Tris • HCl PH 7.5 and 150 mM NaCl, the pH of solution was finally adjusted to around 7.5 followed by DNA duplex anneal (29). We obtained the averaged concentration of protein and DNA samples based on three times measure by Nano Drop ND-1000 spectrophotometer (Thermo Scientific). ITC measurements were performed at the concentrations of MBD domain proteins and DNA oligos ranging from 15 to 50 μM and 0.5 mM to 1 mM, respectively, using the MicroCal ITC or ITC200 (GE Healthcare) at 25 °C. Finally, the dissociation constants (Kds) were determined using Origin 7.0 with one-site binding model (Origin Lab Corp). The standard errors of Kds are the fitting errors from the best ITC titration curves of each binding pair.

### Crystallization of protein-DNA complexes

The purified proteins were mixed with different DNA oligos at a 1:1 molar ratio followed by incubating 30 min on ice. The protein–DNA complexes were crystallized using the sitting drop vapour diffusion method at 18 °C by mixing 0.5 μl of the complex samples with 0.5 μl of the reservoir solution. Finally, we obtained the complex crystals from different conditions. The detailed crystallization conditions were summarized in the Table S1. For data collection, the crystals were then soaked in the reservoir solution containing additional 15% (v/v) glycerol before flash-frozen using liquid nitrogen.

### Data collection and complex structure determination

Diffraction data were collected at beamline 19ID of the Advanced Photon Source and processed with XDS and AIMLESS (30). The MBD3-AmCGT complex structure was solved by molecular replacement with data collected at a rotating anode source on an additional crystal and coordinates from an MBD2-DNA complex. MBD3 complex crystals with GmCGC and AmCGC/GmCGT DNAs, respectively, where virtually isomorphous to the AmCGT complex crystal, obviating renewed molecular replacement searches. The MBD3-GmCGC structure was first refined against lower resolution data from an additional crystal of the MBD3-GmCGC complex.

The MBD1-AmCGT complex structure was solved by molecular replacement with MBD3-AmCGT coordinates. In the course of this structure’s model refinement, a larger free set of reflections was assigned. This reassignment was followed by coordinate randomization with the CCP4 (31) PDBSET NOISE command in order to de-correlate newly free reflections from the model. Molecular replacement searches were performed with the programs PHASER (32) and MOLREP (33). Models were interactively rebuilt in COOT (34), refined with REFMAC (35) and validated with PHENIX.MOLPROBITY (36). Data collection and refinement statistics are summarized in Table S1.

## SUPPLEMENTAL INFORMATION

Supplemental Information can be found online.

## ACCESSION NUMBERS

Coordinates and structure factors for the structures of the MBD1 and MBD3 MBD domains in complex with different DNA ligands have been deposited into Protein Data Bank (PDB) under the accession codes: 6D1T, 6CCG, 6CEU, 6CEV and 6CC8.

## ACKNOWLEDGEMENTS

We thank Wolfram Tempel for structure determination and manuscript reading. We thank Amy Wernimont for reviewing some earlier versions of crystallographic models. Results shown in this report are derived from work performed at Argonne National Laboratory, Structural Biology Center (SBC) at the Advanced Photon Source. SBC-CAT is operated by UChicago Argonne, LLC, for the U.S. Department of Energy, Office of Biological and Environmental Research under contract DE-AC02-06Ch11357. The SGC is a registered charity (number 1097737) that receives funds from AbbVie, Bayer Pharma AG, Boehringer Ingelheim, Canada Foundation for Innovation, Eshelman Institute for Innovation, Genome Canada through Ontario Genomics Institute [OGI-055], Innovative Medicines Initiative (EU/EFPIA) [ULTRA-DD grant no. 115766], Janssen, Merck KGaA, Darmstadt, Germany, MSD, Novartis Pharma AG, Ontario Ministry of Research, Innovation and Science (MRIS), Pfizer, São Paulo Research Foundation-FAPESP, Takeda, and Wellcome. This work is also supported by National Natural Science Foundation of China (31770834 and 31300629).

## AUTHOR CONTRIBUTIONS

K.L. and M.L. purified and crystallized the protein; K.L. and M.L. conducted the ITC assays with assistance from B.G and H.C.; Y.L. cloned some of the MBD domains. K.L. and J.M. wrote the manuscript and all authors contributed to editing the manuscript.

---

**Table S1.**
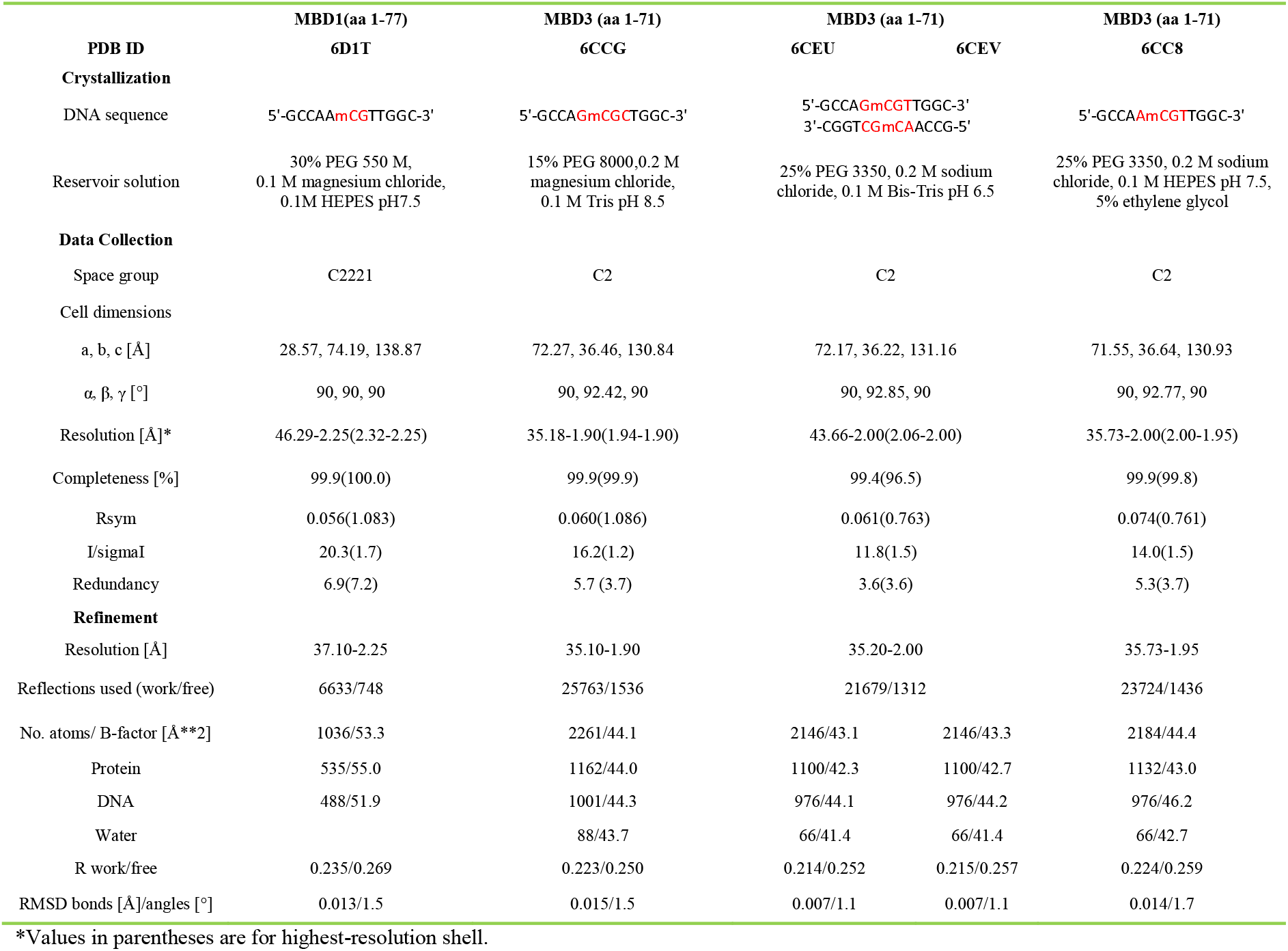
Data collection and refinement statistics

| PDB ID | MBD1(aa 1-77) | MBD3 (aa 1-71) | MBD3 (aa 1-71) |  | MBD3 (aa 1-71) |
| --- | --- | --- | --- | --- | --- |
|  | 6D1T | 6CCG | 6CEU | 6CEV | 6CC8 |
| **Crystallization** |  |  |  |  |  |
| DNA sequence | 5'-GCCAA**mCG**TTGGC-3' | 5'-GCCA**GmCGC**TGGC-3' | 5'-GCCA**GmCGT**TGGC-3'<br>3'-CGGT**CGmCA**ACCG-5' |  | 5'-GCCA**AmCGT**TGGC-3' |
| Reservoir solution | 30% PEG 550 M,<br>0.1 M magnesium chloride,<br>0.1M HEPES pH7.5 | 15% PEG 8000,0.2 M<br>magnesium chloride,<br>0.1 M Tris pH 8.5 | 25% PEG 3350, 0.2 M sodium<br>chloride, 0.1 M Bis-Tris pH 6.5 |  | 25% PEG 3350, 0.2 M sodium<br>chloride, 0.1 M HEPES pH 7.5,<br>5% ethylene glycol |
| **Data Collection** |  |  |  |  |  |
| Space group | C2221 | C2 | C2 |  | C2 |
| Cell dimensions |  |  |  |  |  |
| a, b, c [Å] | 28.57, 74.19, 138.87 | 72.27, 36.46, 130.84 | 72.17, 36.22, 131.16 |  | 71.55, 36.64, 130.93 |
| $\alpha, \beta, \gamma$ [°] | 90, 90, 90 | 90, 92.42, 90 | 90, 92.85, 90 | | 90, 92.77, 90 |
| Resolution [Å]* | 46.29-2.25(2.32-2.25) | 35.18-1.90(1.94-1.90) | 43.66-2.00(2.06-2.00) |  | 35.73-2.00(2.00-1.95) |
| Completeness [%] | 99.9(100.0) | 99.9(99.9) | 99.4(96.5) |  | 99.9(99.8) |
| Rsym | 0.056(1.083) | 0.060(1.086) | 0.061(0.763) |  | 0.074(0.761) |
| I/sigmaI | 20.3(1.7) | 16.2(1.2) | 11.8(1.5) |  | 14.0(1.5) |
| Redundancy | 6.9(7.2) | 5.7 (3.7) | 3.6(3.6) |  | 5.3(3.7) |
| **Refinement** |  |  |  |  |  |
| Resolution [Å] | 37.10-2.25 | 35.10-1.90 | 35.20-2.00 |  | 35.73-1.95 |
| Reflections used (work/free) | 6633/748 | 25763/1536 | 21679/1312 |  | 23724/1436 |
| No. atoms/ B-factor [Å**2] | 1036/53.3 | 2261/44.1 | 2146/43.1 | 2146/43.3 | 2184/44.4 |
| Protein | 535/55.0 | 1162/44.0 | 1100/42.3 | 1100/42.7 | 1132/43.0 |
| DNA | 488/51.9 | 1001/44.3 | 976/44.1 | 976/44.2 | 976/46.2 |
| Water |  | 88/43.7 | 66/41.4 | 66/41.4 | 66/42.7 |
| R work/free | 0.235/0.269 | 0.223/0.250 | 0.214/0.252 | 0.215/0.257 | 0.224/0.259 |
| RMSD bonds [Å]/angles [°] | 0.013/1.5 | 0.015/1.5 | 0.007/1.1 | 0.007/1.1 | 0.014/1.7 |

\*Values in parentheses are for highest-resolution shell.

